# Detecting High Scoring Local Alignments in Pangenome Graphs

**DOI:** 10.1101/2020.09.03.280958

**Authors:** Tizian Schulz, Roland Wittler, Sven Rahmann, Faraz Hach, Jens Stoye

## Abstract

**Motivation:** Increasing amounts of individual genomes sequenced per species motivate the usage of pangenomic approaches. Pangenomes may be represented as graphical structures, e.g. compacted colored de Bruijn graphs, which offer a low memory usage and facilitate reference-free sequence comparisons. While sequence-to-graph mapping to graphical pangenomes has been studied for some time, no local alignment search tool in the vein of BLAST has been proposed yet.

**Results:** We present a new heuristic method to find maximum scoring local alignments of a DNA query sequence to a pangenome represented as a compacted colored de Bruijn graph. Our approach additionally allows a comparison of similarity among sequences within the pangenome. We show that local alignment scores follow an exponential-tail distribution similar to BLAST scores, and we discuss how to estimate its parameters to separate local alignments representing sequence homology from spurious findings. An implementation of our method is presented, and its performance and usability are shown. Our approach scales sublinearly in running time and memory usage with respect to the number of genomes under consideration. This is an advantage over classical methods that do not make use of sequence similarity within the pangenome.

## 1 Introduction

### Motivation

Substantial technological advances in DNA sequencing made genomic data become one of the largest types of information kept by humankind [40]. Thus, finding efficient ways of storing and analyzing this data is of high importance. The discipline of *computational pangenomics* tries to cope with this challenge [29]. A *pangenome* is defined as a set of genomic sequences that may be stored and analysed collectively while being represented as a single entity. Sequences within these sets are usually strongly related and thus highly similar. Commonly, a pangenome comprises all genomic sequences of a species, but this definition may be widened or tightened to any other taxonomic unit or kind of distinction. The pangenomic approach allows a high memory saving potential as sequence parts shared by multiple genomes have to be stored only once. In addition, it enables the simultaneous comparison of a large number of individual genomes while avoiding classical reference-based analyses which turned out to have shortcomings in various cases [6, 10].

Pangenomes can be represented in many forms, ranging from pure collections of raw sequences [43] over alignment based approaches using multiple sequence alignments [14, 34] to graphical structures (e.g. [11, 16, 19]). In this work, we focus on graphical structures. In particular, we represent pangenomes as compacted colored de Bruijn graphs [28, 30]. Advantages of this representation over others include low memory and storage footprints, versatility in accepting different input types (raw sequences or assemblies), and fast and alignment-free construction.

### Background

Basic kinds of comparisons on a pangenome are different variants of sequence searches. Detecting exact or approximate matches can serve to answer questions like membership queries. More complex analyses often involve alignment based methods.

Algorithms for sequence-to-graph alignment have already been studied for some time. Early works date back to 1989 where a graph was used for approximate regular expression mapping [32]. In 2000, an algorithm to align sequences to arbitrary graphs was proposed in the context of hypertext search [33]. Lee *et al*. introduced *partial order alignment* (POA) on directed acyclic graphs in 2002 [25]. Works by Dilthey *et al*. are restricted to acyclic graphs as well [11, 12]. The tool “vg” uses POA on general graphs [16] by *unrolling* cyclic parts. Read mapping on de Bruijn graphs can be done by a heuristic proposed by Limasset *et al*. [26]. Recently, a method was published allowing exact read mapping on general graphs [36]. It is limited to semi-global alignment though. Other solutions have been presented by Antipov *et al*. [5] and Kavya *et al*. [22].

These solutions address a sequence mapping scenario where query sequences are aligned either globally or locally to the graph region that best matches the query. This approach is usually designed for rapidly mapping a huge number of queries gaining speed by the underlying assumption that a query either maps to exactly one position in the graph, or to none.

In this work, we study the problem of finding high scoring local alignments between a query sequence and a graph that are likely to represent sequence homology. The exact notion of “high-scoring” is based on statistical considerations. Hence, we are within the regime of the popular basic local alignment search tool “BLAST” [4]. Even though BLAST is still widely used, many other solutions have been presented for heuristic sequence alignment searches (e.g. [23]). Some exploit new algorithmic techniques or data structures to improve either sensitivity, run time or both (e.g. [13, 39, 15]). Others exclusively focus on protein alignment [46, 41, 7, 42]. Recently, the tool BlastFrost [27] appeared, which enables sequence queries on a pangenome graph. It does not calculate alignments, though.

### Contribution

In comparison to the above mentioned alignment tools that work on collections of individual sequences, our approach stores genomic sequences in a graph to analyse them all in parallel. Moreover, by querying the graph, we are able to compare these sequences not only to the query, but also among each other. This has an advantage over other approaches where an all-against-all postprocessing of results would add a quadratic number of comparisons to their running time. To this end, we introduce the notion of a *quorum* and a *search color set* to allow for customized searches in specific parts of the pangenome. For instance, a user might be interested only in alignments shared by the majority of genomes showing a certain phenotype. Finally, we present first results of alignment statistics with the consideration that all sequences in a pangenome are related and highly similar, which is in contrast to the general initial, underlying independence assumptions. To our knowledge, our approach is unique and has never been studied before.

The remainder of this manuscript is organized as follows. In Sec. 2, we define our model, formally state the problem and describe the algorithmic procedure. Sec. 3 contains use cases of our algorithm and comparisons of its performance. In Sec. 4, we discuss our results. The source code of our implementation, instructions on how to generate samples for our statistical parameter estimation and test data used in this paper are available from https://gitlab.ub.uni-bielefeld.de/gi/plast.

## 2 Methods

### 2.1 Basic definitions

A *string* is a sequence of characters drawn from a finite, non-empty set, called *alphabet*. For a given string *s*, we denote its length by |*s*|, the character at position *i* by *s*[*i*] and the substring starting at position *i* and ending at position *j* by *s*[*i*..*j*]. A string of length *k* is called *k-mer*. For any decomposition *s* = *xy*, the (potentially empty) substrings *x* and *y* are called *prefix* and *suffix* of *s*, respectively.

In this work, we assume that all strings are over the DNA nucleotide alphabet Σ_DNA_ = {`A,C,G,T`}. Then, a *query* is a finite string *q* over Σ_DNA_. A *genome* is a set of strings over Σ_DNA_ that can be many millions of short sequences representing raw data produced by a sequencing machine, or a few long sequences representing chromosomes or contigs of a complete or draft assembly. Each genome is identified with a unique color of the *universal color set U*, and the color is assigned to all strings of the genome to distinguish between sequences from different genomes.

### 2.2 Compacted colored de Bruijn graphs

Let *k* ≥ 2. A *compacted colored de Bruijn graph* of dimension *k* over an alphabet Σ and a color set *U* is a vertex-labeled directed graph *G* = (*V, E*, λ, *C*), where each vertex *v* ∈ *V* is labeled with a sequence λ(*v*) of length at least *k*, each *k*-mer *r* appears in the label of at most one vertex of the graph, there exists an edge (*v, w*) ∈ *E* from vertex *v* to vertex *w* if and only if the (*k* − 1)-length suffix of λ(*v*) equals the (*k*−1)-length prefix of λ(*w*), and *C* assigns a color set *C*(*r*) ⊆ *U* to each *k*-mer *r* that appears in any vertex label. For convenience, we write |*v*| instead of |λ(*v*)| for the length of the label of *v*. In addition, λ′(*v*) denotes the label of *v* except its last *k* − 1 characters, i.e., λ′(*v*):= λ(*v*)[1..(|*v*| − *k* + 1)].

### 2.3 Locations and truncated paths

We represent a *location* in a compacted colored de Bruijn graph *G* = (*V, E*, λ, *C*) by a pair (*v, i*) with *v* ∈ *V* and 1 ≤ *i* ≤ |*v*|. We say that a *k*-mer *r overlaps* a location *l* = (*v, i*) if and only if *r* is a substring of λ(*v*) starting at position *o* such that max(1, *i* − *k* + 1) ≤ *o* ≤ min(*i*, |*v*| − *k* + 1). For an interval [*b*..*e*] of positions within *v*, 1 ≤ *b* ≤ *e* ≤ |*v*|, we define the *interval location* as *L*(*v, b, e*) = {(*v, b*), (*v, b* + 1),…, (*v, e*)}. The interval location *L*(*v*, 1, |*v*|) of a complete vertex *v* is denoted by *L*(*v*).

We say that *p* = (*v*_0_, *v*_1_,…, *v_z_*) is a *path* in *G* if *v*_0_, *v*_1_,…, *v_z_* ∈ *V* and (*v_i_*, *v*_*i*+1_) ∈ *E* for all *i* with 0 ≤ *i* < *z*. A triple *t* = (*p, b, e*) is called a *truncated path* in *G* from location (*v*_0_, *b*) to location (*v_z_, e*) passing vertices *v*_1_,…, *v*_*z*−1_ if and only if (i) *p* is a path in *G*; (ii) *b* ∈ {1,…, |*v*_0_|}; (iii) *e* ∈ {1,…, |*v_z_*|}; and (iv) if *z* = 0 then *b* ≤ *e*. A truncated path *t* = (*p, b, e*) has a sequence

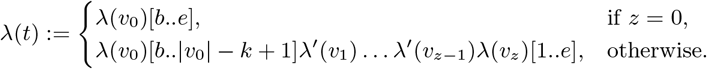

The *path location* of a truncated path *t* = (*p, b, e*) is defined as

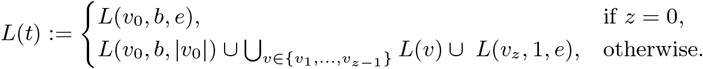

The color set of a path location *L* is then defined as

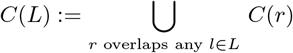

### 2.4 Formal problem statement

Our algorithm finds high scoring local alignments between a given query sequence *q* and a pangenome represented as a compacted colored de Bruijn graph *G* = (*V, E*, λ, *C*) over the DNA alphabet and a color set *U*. Apart from *q* and *G*, it takes as input a non-empty color set *C*_search_ ⊆ *U* called *search color set* and a quorum *m* ∈ {1,…, |*C*_search_|}. It outputs a set 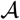, where each 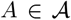 is a pair (*α*, [*c*_1_, *c*_2_,…]) consisting of an alignment *α* between a substring of *q* and the sequence of a truncated path λ(*t*), and a list of checkpoints *c*_1_, *c*_2_,… partitioning *t* into maximal subpaths with the same color set, each containing a subset of *C*_search_ of cardinality at least *m*. Our algorithm allows to find local alignments for both *q* and its reverse complement. For the sake of simplicity, we forgo to mention the reverse complement in the following descriptions. Procedures may be assumed to work similarly for the reverse complementary sequence.

### 2.5 Algorithm

The algorithm consists of three basic algorithmic steps. In the *seed detection step*, maximal exact matches of a minimal length *w* ≤ *k* are searched between *q* and all sequences of *G*. Found seeds are extended without gaps in the *seed extension step*. Statistical considerations are used to extract biologically meaningful findings. These are then recalculated in the *gapped recalculation step* to obtain the finally reported alignments.

#### Preprocessing: graph construction and index generation

While the query sequence *q* has to be specified as an input parameter, the graph *G* may also be constructed from a set of genomes in a preprocessing step. Once *G* has been created for a specific value of *k*, an additional index is built according to a minimal seed length *w* ≤ *k*.

The index consists of two vectors that allow a fast location of exact matches between *q* and sequences stored in *G*. Vector PROF is a *w*-mer profile that contains, in form of a cumulative sum, for any *w*-mer *ω* the number of occurrences of *ω* in *G*, denoted *occ*(*ω*). In other words, PROF[0] = occ(*ω*_0_) and

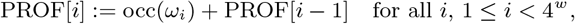

where *ω_i_* is the *i*-th *w*-mer over Σ_DNA_ in lexicographic order. Here, *w*-mers occurring in the (*k* – 1)-length label overlap of two consecutive vertices are counted only once. The second vector, OCC, harbors the occurrences of all *w*-mers appearing in *G*. It stores the location of any seed occurrence, consisting of the ID of a vertex *v* ∈ *V* and the offset in λ(*v*) at which the corresponding *w*-mer starts. All occurrences of a *w*-mer *ω_i_* in *G* may then be accessed by using a ranking function to get position *i* in PROF. PROF[*i* – 1], in turn, gives the offset in OCC where all PROF[*i*] − PROF[*i* – 1] occurrences of *ω_i_* in *G* are stored consecutively. In case a *w*-mer falls into a (*k* – 1)-overlap between two vertices *u* → *v*, the occurrence in *v* is stored. An example of the index and its use can be found in Fig. 1.

**Figure 1:**
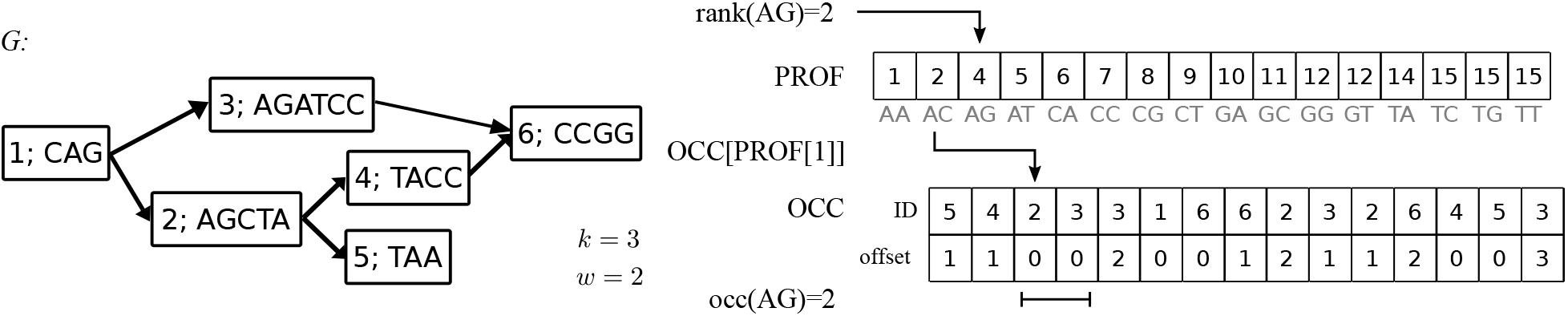
Example of an index search using vectors PROF and OCC. Occurrences of the 2-mer `AG` in graph *G* are accessed by looking up `AG`’s position in PROF which provides the offset in OCC where *occ*(`AG`) locations corresponding to occurrences of `AG` in *G* are stored consecutively. Vector and sequence indices start with 0.

#### Seed detection step

The index is used to look up all *w*-mers occurring in *q*. All matches are checked for quorum fulfillment. Consecutive matches may be merged into a single interval location. A match is stored in a list directly linked to the vertex whose sequence it was found in. This allows a quick access to the match if needed during seed extension. Inside the list, matches are ordered by increasing starting offset in the vertex label. Match order is used to terminate a list iteration as soon as it is clear that a demanded match cannot be part of a list anymore, which leads to an additional speed gain.

#### Seed extension step

Each match represents an interval location used as seed for an ungapped extension. The extension is performed according to an *X*-drop algorithm as used in BLAST along *q* and all truncated paths of *G* starting at an initial interval location (*v,b,e*). The paths are generated by a depth first search traversal through *G* outgoing from vertex *v*. The exploration of a path stops as soon as either the current extension’s score drops below *X* or the quorum requirement is no longer fulfilled. Our experience shows that the exhaustive processing of all existing truncated paths starting from *v* through *G* is possible without a notable slow-down of the algorithm most of the time even though the number of paths may be very large. However, an iteration over all truncated paths can become infeasible if *G* has some dense regions consisting of vertices with short labels and high degrees, producing truncated paths with highly similar sequences. If these sequences are similar to the query *q*, the *X*-drop criterion does not suffice to terminate path explorations leading to suboptimal alignments within an acceptable time frame. Therefore we introduced a threshold that limits the number of vertices which can be visited during the extension of a seed.

#### Gapped recalculation step

The result of the ungapped extension is a set of path locations consisting of a few biologically meaningful and many more spurious hits. Statistical significance criteria, described in Sec. 2.6, are used to rank the hits and separate both kinds. The most significant alignments are recalculated using standard gapped alignment with banded dynamic programming. The band width is chosen according to the quality of the alignment. The final alignments are reported, together with statistics for gapped alignments.

### 2.6 Alignment statistics on a sequence graph

In classical pairwise (linear) sequence comparison, an extensive statistical theory exists, sometimes referred to as Karlin-Altschul theory [21], but subsequently extended, refined and made practical by many others (e.g. [35, 44]). No statistical theory currently exists for sequence queries against (graphical) pangenomes, so we provide a baseline here.

A key question is as follows: Given a score value *s*, how many hits *H_s_* of a random query sequence of length *n* against the given reference (genome or pangenome) of size *m* are observed whose score reach or exceed *s*? The expected value 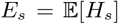 is called the *E-value* of an observed score *s*. Let *S* be the score of the best hit. Then the event {*S* ≥ *s*} is equivalent to {*H_s_* ≥ 1} and the probability 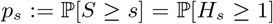 of the event that there exists at least one hit with score reaching *s* is called the *p-value* of an observed score *s*. If *E_s_* is small (say ≤ 0.05) by Poisson approximation and first-order approximation of the exponential function, we have

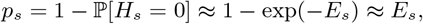

and we need not distinguish in practice between E-value and p-value. We summarize an approximate version of the classical Karlin-Altschul statistics for pairwise sequence alignment and then discuss our proposal about how to generalize the theory for query-to-pangenome alignment.

#### Summary of approximate Karlin-Altschul statistics

Comparing the expected numbers of hits *E_s_*, *E*_*s*+1_, *E*_*s*+2_,… with increasing scores *s*, (*s* + 1), (*s* + 2),…, when already *E_s_* is small, one observes that *E_s+k_* ≈ *μ^k^* · *E_s_* for some factor 0 < *μ* < 1, mostly independent of *s*, as long as *s* is large enough. Factor *μ* is typically written as e^−λ^ with some λ > 0. This holds if the average score when comparing two single nucleotides is negative; otherwise there exist arbitrarily long high scoring matches. If we increase the length *n* of the query or length *m* of the reference, we provide more possibilities for a hit, and the expected number of such hits increases linearly with both *n* and *m*. This leads to

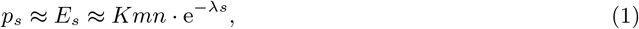

where *K* > 0 and λ > 0 are constants, *n* is the query length, and *m* is the reference genome length. Note that this approximation is only valid for the extreme tail of the distribution (large *s*, small *E_s_* ≈ *p_s_* ≤ 0.05). The functional form of (1) has been empirically shown to be very robust, valid for ungapped and gapped alignments and even when considering compositional bias (difference in GC content) between query and reference, or when considering a fixed query against a random reference [45]. Constants *K* and λ depend on the scoring scheme, including the gap costs for gapped alignment. In practice, λ must be determined by sampling and simulation [3, 45].

#### Statistics for pangenome alignment

We hypothesize that a relation as in Eq. (1) can be observed when considering the top score of a random query aligned against a pangenome, if s is sufficiently large such that hits reaching score s are rare. We further hypothesize that such a relation holds for both ungapped high-scoring pairs after seed extension and for final gapped alignments, albeit with different values of λ > 0 and *K* > 0. However, the dependency of λ and *K* on sequence relatedness and diversity within a pangenome may be complex, and it is out of scope of this work to investigate the details. Instead, we investigate whether indeed there holds an affine-linear relationship log *p_s_* ≈ *C* − λ*s* with 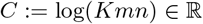 for fixed query length *n* and a pangenome graph of size *m*.

#### Parameter estimation by importance sampling

To obtain a good estimate of λ > 0 and 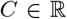, we need good estimates of small probabilities *p_s_* for large *s*. Using random sequences, large *s* with small *p_s_* < 10^−6^ are by definition rare, so too many samples would be needed for accurate estimates. Hence we only use this “naive” random sampling strategy to obtain an initial estimate of *C* and λ and then resort to importance sampling, using a Metropolis-Hastings Markov Chain Monte Carlo (MCMC) strategy similar to the one described by Wolfsheimer *et al*. [45].

In brief, let 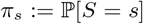 be the unknown score distribution on integers *s*. We construct a Markov chain in such a way that the probability to sample a random sequence with score *s* is exponentially biased towards higher scores, *r_s_*:= *π_s_* · exp(λ_0_ · *s*)/*Z*, where λ_0_ < λ should slightly underestimate the true λ and can be derived from the initial naive sampling step, and *Z* is the appropriate (unknown) normalization constant such that Σ_*s*_ *r_s_* = 1. In order to avoid computing *Z* explicitly, we use the Metropolis-Hastings method: Given a current DNA sequence *x*, a new candidate sequence *y* is proposed from a neighborhood of *x*; see [45] for the precise definition of the neighborhood. Roughly, a single nucleotide can be inserted, deleted or substituted at any position, deleting or inserting a nucleotide at the left or right end to keep the sequence length *n* constant. Thus the edit distance between *x* and the new proposal *y* is at most 2. This creates a Markov chain where, in equilibrium, each sequence is equally probable, similarly to the naive simulation, where each nucleotide is drawn independently from a uniform distribution. Now, to bias the samples toward higher scores, the scores *s_x_* and *s_y_* of *x* and proposal *y*, respectively, are compared. We accept *y* with probability min{1, (*r_s_y__* /*π_s_y__*)/(*r_s_x__*/*π_s_x__*)} = min{1, exp(λ_0_ · (*s_y_* – *s_x_*))}, i.e., a score increase is always accepted, and a decrease only with a certain probability. When a proposal is rejected, *x* stays the current sequence and another proposal is generated. A score sample is drawn after a large number of accepts that allows the query sequence to change considerably in comparison to the previous sample. Typically, 2*n*/3 accepts suffice for a query of length *n*. The first few samples are discarded to allow the Markov chain to reach equilibrium. Since the defined chain is rapidly mixing (there exist short paths from every sequence to every other sequence), we found it sufficient to discard the first 5 samples.

The procedure yields uncorrelated score samples, which stem from the biased distribution *r* = (*r_s_*). Let *R_s_* be the absolute number of times that score *s* was sampled. Let 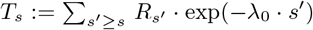. Then λ can be estimated by fitting a line to points (*s*, log *T_s_*) in an interval of *s* where the counts *R_s_* are consistently high, say *R_s_* ≥ 50. Then, *C* is estimated from the 10% of highest scores in the initial naive sampling step.

## 3 Results

We implemented the algorithm described in Sec. 2.5 in C++ using Bifrost [17] as the underlying realization of a compacted colored de Bruijn graph. Among several other implementations (e.g. [2, 8, 18, 19, 31]), we chose Bifrost since it is an efficient, easy-to-use implementation, allows the usage of assembled and raw sequencing data, and provides the possibility to assign any kind of data to vertices of the graph. Our implementation called PLAST (“Pangenome Local Alignment Search Tool”) is available from https://gitlab.ub.uni-bielefeld.de/gi/plast. We present results of our statistical parameter estimation, followed by a performance analysis of our method. In particular, we show the advantage in runtime, memory usage and result aggregation when searching local alignments inside a pangenome with our method compared to a conventional search and analysis using other BLAST-like software tools. Afterwards, we present a more advanced use case and show that our method scales even to human data. Unless stated differently, all graphs have been built for a *k*-mer length of 31 (Bifrost default), and searches were performed using default parameters. To obtain statistical parameters, we used a combination of initial naive sampling followed by importance sampling, as described in Sec. 2.6.

### 3.1 Statistical parameter estimation

We tested the hypothesis from Sec. 2.6 that log *p_s_* ≈ *C* − λ*s* for constants 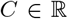 and λ > 0 for both ungapped and gapped alignments. After initial confirming results on simulated pangenomes (not shown), we considered a real pangenome of 220 *Salmonella enterica* genomes of the same lineage (Para C) taken from [47]. Naive simulation was performed with one million random DNA sequences of length *n* = 200, yielding the empirical complementary cumulative distribution function (ccdf) of the best hit’s score for each query for both the gapped and ungapped case. A least-squares fit of affine linear functions in the distribution’s tail, considering logarithmic cumulative relative frequencies between 10^−2^ and 10^−1^, yielded initial estimates of *C* = 17.45 and λ = 1.085 for ungapped alignments and *C* = 14.81 and λ = 0.852 for the final gapped alignments; see Fig. 2. As expected, λ_gapped_ < λ_ungapped_, as gaps provide more freedom to achieve a higher score with the same query length. The affine relationship cannot hold for much higher p-values because our approximation assumes small *p_s_* ≤ 10^−2^, and we also cannot make a statement for much lower p-values with “only” 10^6^ simulations.

**Figure 2:**
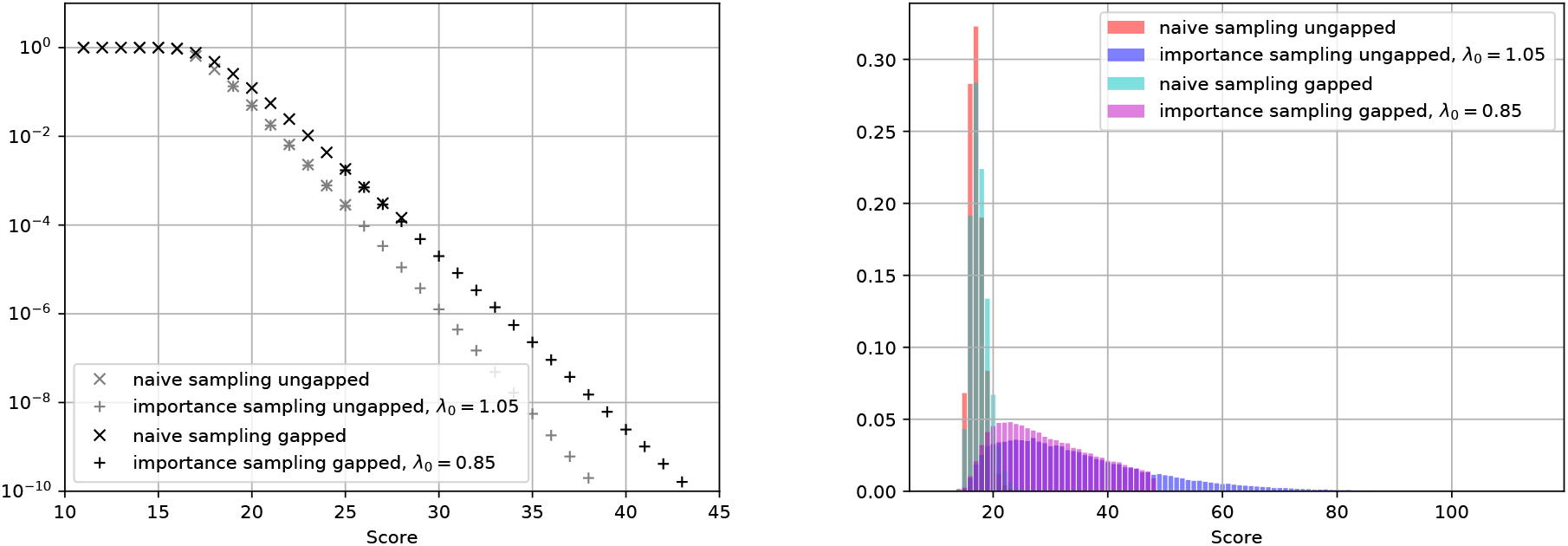
Results of naive and importance sampling. **Left:** Logarithmic plot of p-values (complementary cumulative distribution function) of highest ungapped and gapped alignment scores for random queries (without color or quorum constraints) against a pangenome of 220 *Salmonella enterica* genomes. Naive simulation with 10^6^ samples yields accurate estimates for p-values in the range between 10^−2^ and 10^−4^. Importance sampling enables a better view of the rare-event tail. For p-values ≤ 10^−2^, the hypothesis of an affine dependency holds with values *C* ≈ 20.78 and λ ≈ 1.136 for ungapped alignments and *C* ≈ 16.11 and λ ≈ 0.898 for gapped alignments, estimated from importance sampling. **Right:** Histograms of (normalized) sample counts per score value for ungapped and gapped alignments, using naive sampling and importance sampling. Importance sampling gives access to a broader interval of scores in the rare-event tails.

To gain access to the rare-event tail, we performed importance sampling as described above. Per sampled sequence and score, we need to evaluate (2*n*/3)/*α* sequences, where *α* is the average acceptance rate, which we typically find to be around 0.46 to 0.75, which amounts to approximately (2*n*/3)/(1/2) = 4*n*/3 evaluated sequences per drawn sample. Thus, importance sampling introduces a 250-fold overhead over naive sampling for *n* = 200. However, it allows us to sample from high scores that are unobtainable by naive sampling, even if billions of samples were used, yielding much higher efficiency by several orders of magnitude (cf. importance sampling tails in Fig. 2). We obtain refined estimates of λ = 1.136 (*C* = 20.78) for ungapped and λ = 0.898 (*C* = 16.11) for gapped alignments. As expected, using weight factors of λ_0_ = 0.852 for gapped and λ_0_ = 1.05 for ungapped that are slightly smaller than the “true” λ = 0.898 for gapped and λ = 1.136 for ungapped, we observe an almost flat and slowly decreasing histogram of score counts (Fig. 2) and thus sample from a broad interval of scores.

### 3.2 Comparison to other tools

We evaluated our approach by comparing our implementation against MMseqs2 (as of 08 January 2020), blastn (2.6.0+), BLAT (36×4) and UBLAST (11.0.667). The tools DIAMOND, SWORD and GHOSTZ do not support DNA to DNA alignment. RAPSearch2 was not possible to install due to a reported, but unresolved issue. LAST is known to run very slow on highly redundant data sets—index building was terminated after 10 days. BLAT was designed to search genomes (represented as target databases) for query sequences. It was only possible to run the tool for 750 genomes at once. Runs for larger pangenome sizes had to be split into separate program calls. Results and runtimes were aggregated. Similarly, UBLAST’s freely available 32-bit version has an upper database size limit such that pangenomes consisting of more than 100 genomes had to be distributed on multiple databases.

For the comparison, we downloaded 5,000 randomly selected *Salmonella Typhimurium* assemblies from a total of 19,237 which were annotated as serovar “Typhimurium” from EnteroBase [1]. As queries we chose 100 random substrings of length 1,000 from the *Salmonella* reference genome assembly (RefSeq assembly accession GCF_000195995.1). Our aim was to generate a heterogeneous set of queries fully or partially representing sequences of varying levels of homology.

If possible, tools were run with the same scores for match, mismatch, gaps and the same *X*-drop value. Additionally, we set the maximum number of reported results high enough to get all existing results. This was necessary, because all other tools do not compare their results internally and would otherwise report only the best result for each genome separately, hiding all further findings. For all remaining parameters, default values were used. Concrete program calls are documented at https://gitlab.ub.uni-bielefeld.de/gi/plast. Calculations were performed single threaded on a virtual machine with 28 cores and 256 GB of RAM.

In order to compare the results, we scanned the output of each tool and identified corresponding alignments of PLAST and BLAST. Two results are considered *matching* if they overlap by at least 90 % of the shorter alignment. We call an alignment *unique* if it does not match any alignment of another tool. The comparison is shown in Tab. 1. We see that PLAST reports the lowest total number of results. This is due to the fact that one alignment of PLAST may correspond to several genomes of the pangenome. All other tools report such alignments separately, and an additional postprocessing step would be required to merge them. UBLAST reports by far the highest number of alignments. Thus, it seems to be most sensitive in our experiment. Apart from UBLAST, the number of results which could be found by some tool but not PLAST is below 0.03% (columns “Tool\PLAST” and “BLAST\Tool”, first row). The percentage of results unique to PLAST varies between ~4% and ~6% among the tools. However, only 178 PLAST alignments (2.4%) were unique w.r.t. *all* other tools. Their score was low (mean 18.5, maximum 25) and they were short (mean length 28bp, maximum length 112 bp). They are a result of the different statistical parameters used by PLAST in comparison to other tools. As explained in Sec. 1, these tools do not consider a pangenomic use case where all sequences in the database are highly similar. Thus, they overestimate the chance for a random hit. Using the same statistical parameters as BLAST, all 178 results are filtered out by PLAST due to the E-value threshold. We observed 306 PLAST alignments (4.0%) which were unique w.r.t. at least one other tool but at the same time found by at least one other. 56 of them had a score higher than 25 (maximum 678) or were longer than 112 bp (maximum 999 bp).

**Table 1:** Comparison of PLAST to other alignment tools. 100 random substrings of length 1,000 from the *Salmonella* reference genome assembly have been aligned to the “Typhimurium” dataset. The columns list the tool names, absolute number of alignments, and a comparison to PLAST and BLAST, where X\Y denotes the number of results of X that did not match any result of Y.

| Tool | Results | Tool\PLAST | PLAST\Tool | Tool\BLAST | BLAST\Tool |
| --- | --- | --- | --- | --- | --- |
| PLAST | 7,565 | – | – | 357<br>4.72 % | 290<br>0.02 % |
| BLAST | 1,246,221 | 290<br>0.02 % | 357<br>4.72 % | – | – |
| BLAT | 457,089 | 1<br>0.00 % | 456<br>6.03 % | 508<br>0.11 % | 49,798<br>4.00 % |
| MMseqs2 | 695,792 | 6<br>0.00 % | 322<br>4.26 % | 800<br>0.12 % | 21,022<br>1.69 % |
| UBLAST | 4,881,509 | 111,386<br>2.28 % | 272<br>3.59 % | 220,577<br>4.52 % | 5,459<br>0.44 % |

Next, we randomly selected subsamples from the 5,000 *Salmonella* genome assemblies to generate pangenomes of different sizes and compared the tools’ performance. Plots showing runtime and memory usage may be found as supplemental material.Generally, we can see that most tools quickly lose speed with a growing pangenome size. Only MMseqs2 is able to keep a speed comparable to PLAST even for a pangenome of 5000 individual genomes. However, its speed comes along with a huge memory requirement which is far beyond that of all remaining tools.

All tools spend a large amount of memory for loading the highly redundant data sets and additional index data structures for fast alignment searches. Our graphical representation, in turn, allows a maximal exploitation of sequence similarity. Most parts of the graph represent sequences shared by many individual genomes that have to be stored only once. An alignment calculated for these parts is valid for all sharing genomes and does not have to be calculated multiple times. Thus, we observe a strongly reduced growth in run time and memory consumption for PLAST. Considering a pangenomic use case with constantly growing numbers of individual genomes, our approach will always be superior compared to conventional methods not making use of sequence similarity.

When comparing PLAST’s run time for different quorum values and pangenome sizes, we can generally observe two driving forces (data not shown). On the one hand, the usage of a quorum has a beneficial influence on PLAST’s run time, because it allows to disregard parts of the graph harboring rarely appearing variations which may prune it considerably, especially if the overall diversity within the pangenome is high. At the same time, a quorum check is comparatively cheap if the graph has only a moderate total number of colors in it. On the other hand, for larger pangenomes, the usage of a high quorum becomes an increasing burden as color coverages have to be checked on every vertex using Bifrost’s API. The high degree of color compression generates a noticeable loss in speed in this case which outstrips any gain by pruning from a certain pangenome size. In our experiments, it doubled computation time compared to a run without quorum for the largest pangenome size.

In order to prevent this loss in speed, a first filter was implemented that, instead of only checking color presence, also considers the number of missing colors on a vertex and rejects it as soon as this number becomes too high during iteration. Other heuristics to speed up a quorum check are currently under development and discussed in Sec. 4.

### 3.3 Pathogenicity islands in *Vibrio cholerae*

Pandemic strains in *Vibrio cholerae* are known to contain the *Vibrio Pathogenicity Island-1* (VPI-1) that consists of a whole set of genes whose sequence and order inside the island can be diverse among different strains [20].

We examined a recent collection of 21 *Vibrio cholerae* genomes, 7 of which have been obtained from clinical samples and are labelled “pandemic genomes” (PG), and the remaining 14 have been obtained from non-clinical samples and are labelled “environmental genomes” (EG) [37, primary dataset]. Some of the samples were available as assemblies whereas others could be found only as raw read datasets. Nevertheless, one graph was built from all 21 samples, using the functionality of the underlying Bifrost library to construct a de Bruijn graph from both types of data: read data, that is automatically filtered for low coverage *k*-mers, as well as assembly data that is not filtered.

As a proof of concept, we used PLAST to search VPI-1 inside the *Vibrio cholerae* pangenome - once restricting the search to alignments with PG genomes to verify the existence of VPI-1 in PG, and once restricting the search to alignments with EG genomes to verify the absence of VPI-1 in EG. For the search, we set an E-value cut-off of 0.01, the maximum number of alignments to maximum, and used standard or automatically determined parameters otherwise.

The order of VPI genes within PG genomes may be rearranged, and EG genomes might contain only small fragments of the island. Even so, we used the complete island sequence (accession no. AF325734 [20]) of length 41,272,bp as query. PLAST’s ability to compute *local* alignments and to report many sub-optimal findings, nevertheless allowed an easy detection of each VPI gene separately. Furthermore, conserved gene orders could be detected by local alignments spanning larger fragments of the island. We observed 64 alignments in the PG search that were longer than 2,814 bp, i.e., three times the median length of coding sequences in VPI-1.

For each coding sequence in VPI-1 we determined the maximum overlap by any local alignment. As can be seen in Fig. 3, when restricting the alignment to PG, all coding sequences are covered by alignments almost completely (row “PG”). In contrast, when restricting the alignment to EG, only very few (three of twenty-nine) coding sequences are covered by any alignment by 50 percent or more (row “EG”). We want to highlight here that the detection of such outstanding sequences that are contained in any of a whole group of genomes is possible by a single PLAST search. For further investigations, it would be possible to analyze the PLAST alignments for individual genomes, as exemplified in the remaining rows in Fig. 3, where the maximum overlap is determined among those parts of the alignments that are supported by the corresponding individual genomes.

**Figure 3:**
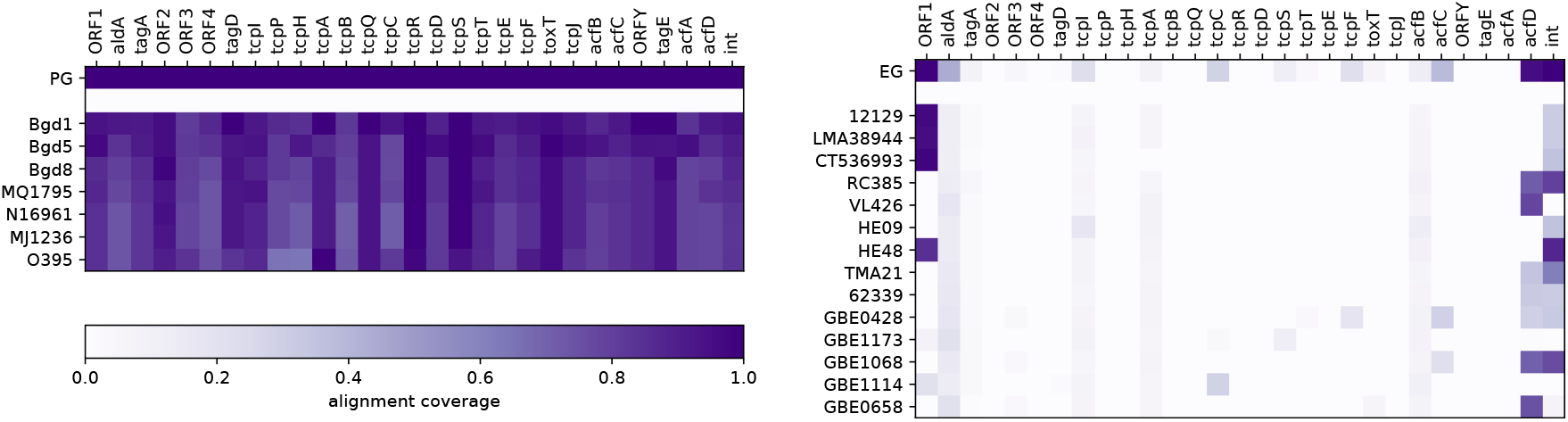
PLAST search results for VPI-1 [20] inside the *Vibrio cholerae* dataset [37] with search color set PG (left) and EG (right). Columns correspond to coding sequences in VPI-1 ordered as appearing in the query sequence. Rows “PG” and “EG” show the maximum overlap of any alignment with a coding sequence inside the pangenome using PG or EG as search color set, respectively. Rows “Bgd1” to “O395” and “12129” to “GBE0658” show the maximum coverage of a sample in any alignment overlapping a coding sequence using PG or EG as search color set, respectively.

### 3.4 Beyond bacterial pangenomes

In order to test PLAST’s applicability beyond bacterial pangenomes, we built a human pangenome using data from the 1000 Genomes (1KG) Project phase 3 [9]. In particular, we used bcftools consensus (https://github.com/samtools/bcftools) to generate chromosome-wise genomic sequences of all 2,504 human individuals by inserting all reported variations into the GRCh37 reference sequence and built graphs (*k* = 63) for chromosomes 2 and 15.

We exemplarily investigated a known polymorphism within the human pangenome. The single nucleotide polymorphism *rs1426654* is reported to influence skin pigmentation. Its reference allele indicates a light skin color which is common to a West Eurasian ancestry [24, 38]. It is located on exon 3 of gene *SLC24A5* on chromosome 15. Searching the exon sequence from reference genome GRCh37 within the pangenome of chromosome 15 resulted in two full size alignments: a perfect match representing the reference allele, and one having a single mismatch representing the variant allele. Restricting our search to the European core genome (search set of all European samples and a quorum of 99%), we exclusively found the reference allele, confirming the results of the earlier studies.

To evaluate PLAST’s performance on the human pangenome, we used a graph of chromosome 2 (≈ 8% of the complete human genome) and searched 100 queries of length 1,000 randomly drawn from the human reference genome. For a comparison, we also ran MMseqs2, next to PLAST the fastest tool according to Sec. 3.2, on the same queries. PLAST took 317s per query on average for the complete data set using a maximum of 24 GB of memory and running on a single thread. Only using a subset of 1,000 chromosomes, MMseqs2 took 339s per query on average using a maximum of 259 GB of memory and running on all available 28 cores. Furthermore, its input files (sequences and index structures) required 2.2 TB of disk space. PLAST’s input files occupied only 9.2 GB on disk in total. Both tools were run with default parameters.

## 4 Discussion

We presented a new BLAST-like method to find highest scoring local alignments between a query sequence and all sequences of a pangenome represented as a colored de Bruijn graph. Unlike read mapping tools developed to find the best or possibly a few suboptimal mapping positions for potentially many query sequences in a short time, our aim is to find all such alignments with statistically significant score. Using a minimal seed length much smaller than *k* increases our detection sensitivity which goes far beyond the scope of a *k*-mer-based seeding approach, while alignment statistics enable us to filter the results for biologically meaningful ones.

By working on a graph, our method is able to exploit the high degrees of sequence similarity within a pangenome. On one hand, this allows to draw conclusions not only about the similarity towards a query sequence, but also to compare the diversity of genomic sequences in the graph with regard to the query. On the other hand, it avoids the storage and processing of highly redundant information and lets the run time and memory usage of our algorithm scale sublinearly with respect to the number of genomes inside the pangenome. Both advantages make our approach superior in comparison to conventional local alignment search tools working on a database of multiple individual genomes, scaling linearly in the number of sequences and enables us to even handle large eukaryotic pangenomes. We showed this in a comparison to other state-of-the-art BLAST-like alignment search tools and by using human data from the 1KG Project.

Moreover, we introduced the usage of a quorum and a search color set which allow to limit searches on customized regions of the pangenome. This is extremely useful for answering specific research questions. Additionally, it avoids repeated construction of the graph for different database sequence sets, which is especially important if pangenomes are large and graph construction becomes prohibitively expensive. As a practical application of our algorithm, we demonstrated its usability on a classical pangenomic use case in Sec. 3.3.

Although our implementation PLAST performs well in practice, it has to be seen as a proof of concept implementation so far. For example, no affine gap cost model has been incorporated yet and is subject to ongoing work. Also, many heuristics are strongly oriented on common techniques for plain sequences. We are convinced that further efforts on the development of heuristics exploiting the special conditions prevailing in a graphical pangenome may lead to even more efficient algorithms.

Yet, quorum checks may become a time determining factor if large pangenomes are considered and quorums are high. The reason is that Bifrost compresses color information within binary matrices for each vertex, and accessing this information requires a time consuming iteration over the matrix. Currently, we are working on ways to aggregate this information as a preprocessing step next to index building to avoid matrix iterations during the search. Aggregated quorum information could be stored for each vertex, e.g., by using five integers encoding the minimum quorum fulfilled on the vertex, the potentially higher quorum being fulfilled at its beginning and at the end, and the sequence offsets at which this higher quorum breaks. A different idea would be to store the number of colors covering a *k*-mer separately for each *k*-mer using a single byte. The memory footprint for both ideas would be small especially in pangenomes of very closely related individual genomes.

We established that, similarly to the statistical behavior of pairwise alignment, sequence-to-graph alignment p-values exhibit exponential tails, log *p_s_* ≈ *C* − λ · *s* for constants 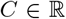, λ > 0. Still, many interesting open questions remain about these statistics. The calculation of precise statistical parameters for each pangenome graph is based on compute-intensive simulations so far. In order to benefit from existing simulations on similar graphs, instead of starting a new simulation for every pangenome graph, we would like to better understand how the statistical parameters change with graph properties and scores. While the dependency on gap scores has been explored in the past, nothing is known about how (and which) graph properties influence the parameters, and preliminary experiments show complex patterns, which we intend to investigate in future work.

We also plan to further extend the concept of quorum and search color set. In particular, we would like to give more freedom to the user by allowing not only to focus on certain parts of the graph, but even to explicitly exclude regions from analysis.

## Acknowledgements

This work was supported by the BMBF-funded de.NBI Cloud within the German Network for Bioinformatics Infrastructure (de.NBI) [031A537B, 031A533A, 031A538A, 031A533B, 031A535A, 031A537C, 031A534A, 031A532B].

## A Appendix

### A.1 Runtime and memory usage plots of tool comparison

**Figure 4:**
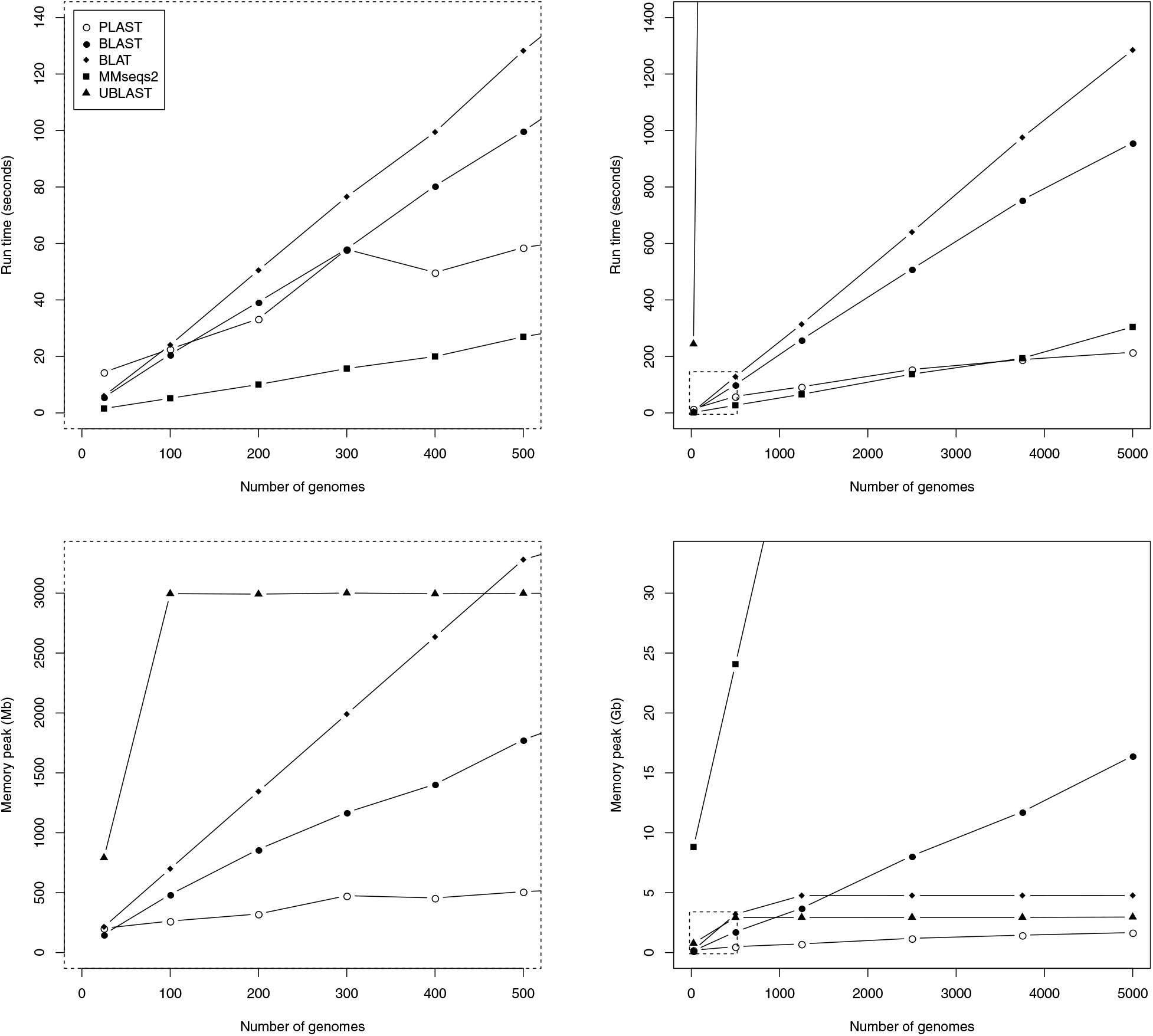
Run time and memory usage comparison of all tools. The areas marked by a dashed rectangle in the plots on the right are shown in separate plots on the left. Values on less than 1000 genomes have been averaged over five random subsamples each. The same legend applies to all plots.

